# Insulin-Like Growth Factor I Couples Metabolism With Circadian Activity Through Hypothalamic Orexin Neurons

**DOI:** 10.1101/2021.02.07.429949

**Authors:** J. Pignatelli, E. Fernandez de Sevilla, J. Sperber, J.D. Horrillo, G. Medina-Gomez, I. Torres Aleman

## Abstract

Uncoupling of metabolism and circadian activity is associated with an increased risk of various pathologies, including neurodegeneration. Recently, insulin and the closely related insulin-like growth factor I (IGF-I) were shown to entrain feeding patterns with circadian rhythms. Moreover, both hormones act centrally to modulate peripheral glucose metabolism; however, whereas central targets of insulin actions are intensely scrutinized, those mediating the actions of IGF-I remain undefined. We analyzed whether IGF-I targets orexin neurons in the lateral hypothalamus, as these neurons are involved in circadian rhythms and energy allocation, and are modulated by IGF-I. Mice with disrupted IGF-IR activity in orexin neurons show phase shifts in circadian feeding behavior, loss of circadian orexin expression, and gradually develop sex-dependent metabolic alterations. In addition, modulation by IGF-I of hepatic KLF transcription factors involved in peripheral glucose metabolism is mediated by orexin neurons. Thus, IGF-I entrains energy metabolism and circadian rhythms through hypothalamic orexin neurons.

## Introduction

In mammals, sleep/wake and food intake cycles are tightly regulated by circadian and metabolic rhythms^1–4^. Circadian rhythms are driven by the hypothalamic suprachiasmatic nucleus (SCN), which is considered the pace-maker region, and other hypothalamic and brainstem regions^4–6^. Metabolic rhythms are driven by clock genes from secondary brain regions, mainly located in the hypothalamus, and peripheral organs such as the liver^4,7^. Circadian rhythms are reset every 24h by environmental light cycles detected by the retina and transmitted to the SCN^6^, while metabolic rhythms are reset by food intake, which is termed the “food clock”^8^. Light entrains the circadian clock through the SCN, and this synchronizes clock genes from peripheral organs through the hypothalamic-pituitary-adrenal axis^9^. In turn, metabolic clocks tune circadian clocks through hormonal (ghrelin, leptin, insulin, glucocorticoids, GLP1) and nutrient (glucose, ketone bodies, non-esterified lipids) levels, and by food-anticipatory behavior^10,11^.

Hypothalamic orexin neurons are important modulators of circadian rhythms such as the sleep/wake cycle^12–14^, and also influence SCN clock cell activity and time-keeping^15^. Insulin-like growth factor I (IGF-I) has recently been postulated to entrain circadian and feeding rhythms at still undefined central sites^16–18^. Moreover, it has long been described that serum IGF-I levels are related to glucose metabolism and insulin sensitivity ^19–21^, putatively acting at a central site^22^. In turn, hypothalamic orexin neurons are involved in glucose sensing^23^ and metabolism^23^, feeding behavior^25^, and overall energy balance^25^, and their impairment are linked to obesity^26^. Orexin neurons are directly modulated by IGF-I ^27^ and diverse humoral signals, including glucose and insulin^28^.

Based on these observations, we postulated that IGF-I acts centrally to entrain hepatic metabolic rhythms with circadian rhythms by modulating the activity of orexin neurons, and in this way participate in the regulation of peripheral glucose metabolism. We now present evidence that IGF-I signaling through orexin neurons is required for glucose homeostasis and proper entrainment of feeding and activity patterns.

## Results

### IGF-I signaling through orexin neurons modulates circadian glucose fluctuations and feeding patterns

To analyze the role of IGF-I on orexin neurons we inactivated its receptor (IGF-IR) in these neurons (Firoc mice), as described^27^. Since we previously observed that Firoc mice have reduced levels of hypothalamic orexin^27^ and levels of this neuropeptide shows a circadian rhythm^29^, we determined whether its circadian fluctuations were also altered in Firoc mice. We measured mRNA levels at ZT8 (inactive phase) and ZT12 (the beginning of the active phase) and found a significant attenuation of the orexin peak at the latter time (Figure 1A). Since both orexin^30^ and IGF-I^22,31^ regulate glucose homeostasis, which also follows a circadian rhythm^5^, we examined levels of circulating glucose in Firoc mice at different time points during the 24 h period and found sex-dependent perturbations. Thus, in Firoc males, blood glucose peaked at ZT8 instead of at Z14 (Figure 1B). In females, glucose levels remained normal, but the peak at Z14 was significantly larger than in littermate females (Figure 1C).

**Figure 1:**
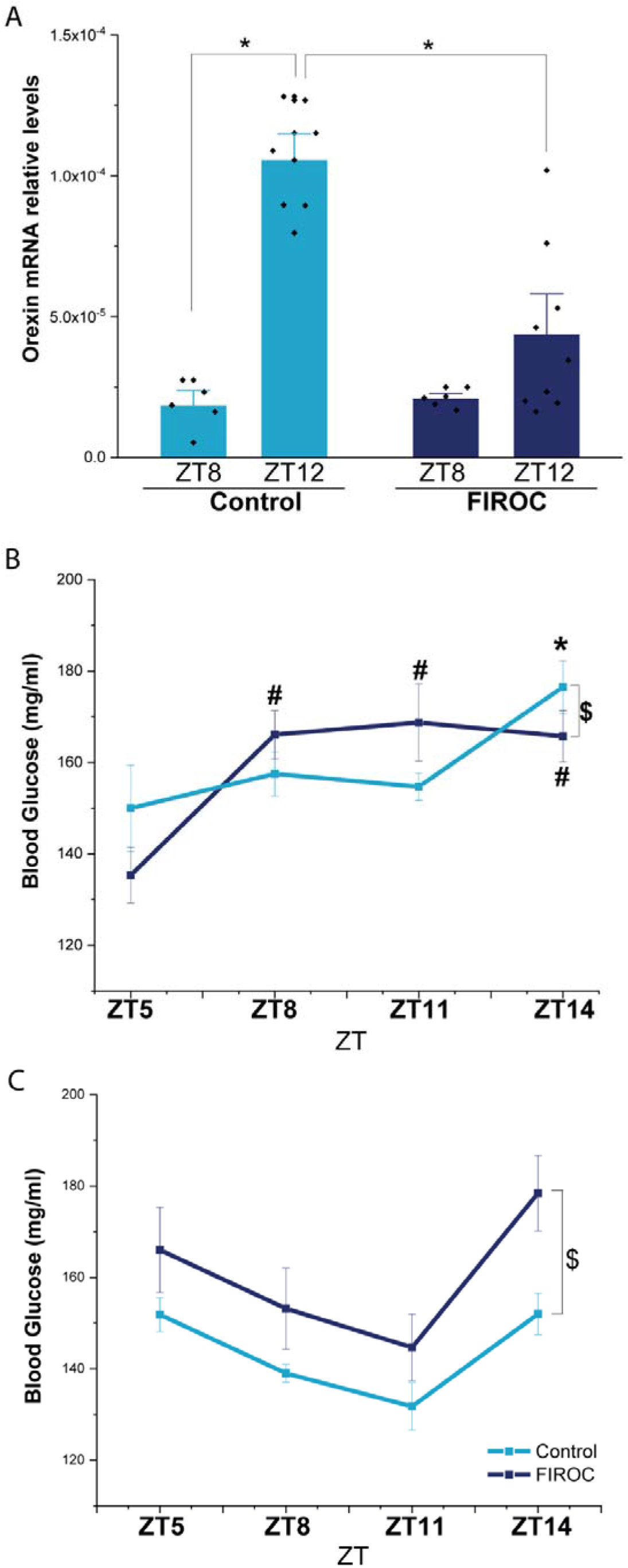
Disruption of IGF-IR in orexin neurons alters orexin and glucose circadian patterns. **A,** Orexin mRNA levels in the hypothalamus of Firoc mice and littermates (Control). Orexin expression peaks in control mice at the end of the resting phase (ZT13), whereas in Firoc mice this peak is significantly reduced (*p<0.05 vs respective controls). **B, C,** Blood glucose was measured at ZT5, ZT8, ZT11, and ZT14 in male (n= 10-15) and female (n=7-10) Firoc mice and littermates. Males showed an advanced rising in glucose levels as compared to controls. Firoc females showed a glucose rhythm similar to controls, but with a significantly higher peak at Z14 (# and *p<0.05 vs ZT5; $p<0.05 vs littermates, t-test).

Since orexin neurons regulate feeding behavior and energy homeostasis^32^, we placed Firoc mice in metabolic cages to analyze feeding and metabolic patterns using an algorithmic analysis to characterize the circadian components of each parameter (Table S1). Although no differences in overall food intake or activity were seen between Firoc mice and littermates, changes in behavioral patterns were observed in a sex-dependent fashion. Male Firoc mice showed significantly increased feeding at ZT12 (Figure 2A), whereas in littermates peak feeding was at ZT15. Thus, a phase shift in feeding rhythm was present (Figure 2A, bottom panel). Consequently, phase advances of about 0.5-1h in respiratory exchange ratio (RER) and energy expenditure (EE) were observed (Table S1). Male Firoc mice had an RER phase of 15.85 ± 0.49h, whereas control littermates had a phase of 16.12 ± 0.99h. Besides, Firoc males had an EE phase of 15.62 ± 0.27h and controls of 16.31 ± 0.75h. No differences were observed in activity in male Firoc mice (Figure 2B). Conversely, Firoc females did not show differences in feeding behavior (Figure 2C), but developed a larger disruption of activity rhythms with a significant decrease at ZT12 (Figure 2D). These changes were reflected in phase shifts of 1-2h; (15.54 ± 0.4h in Firoc, and 18.2 ± 1.5h in controls, Table S1). The phase shift in activity was reflected in metabolic rhythms. Female Firoc mice show an RER phase of 16.50 ± 3.08h whereas littermates had an RER phase of 14.96 ±1.3h, and an EE phase of 15.60 ± 1.23h vs 16.07 ± 1.26h in controls (Table S1).

**Figure 2:**
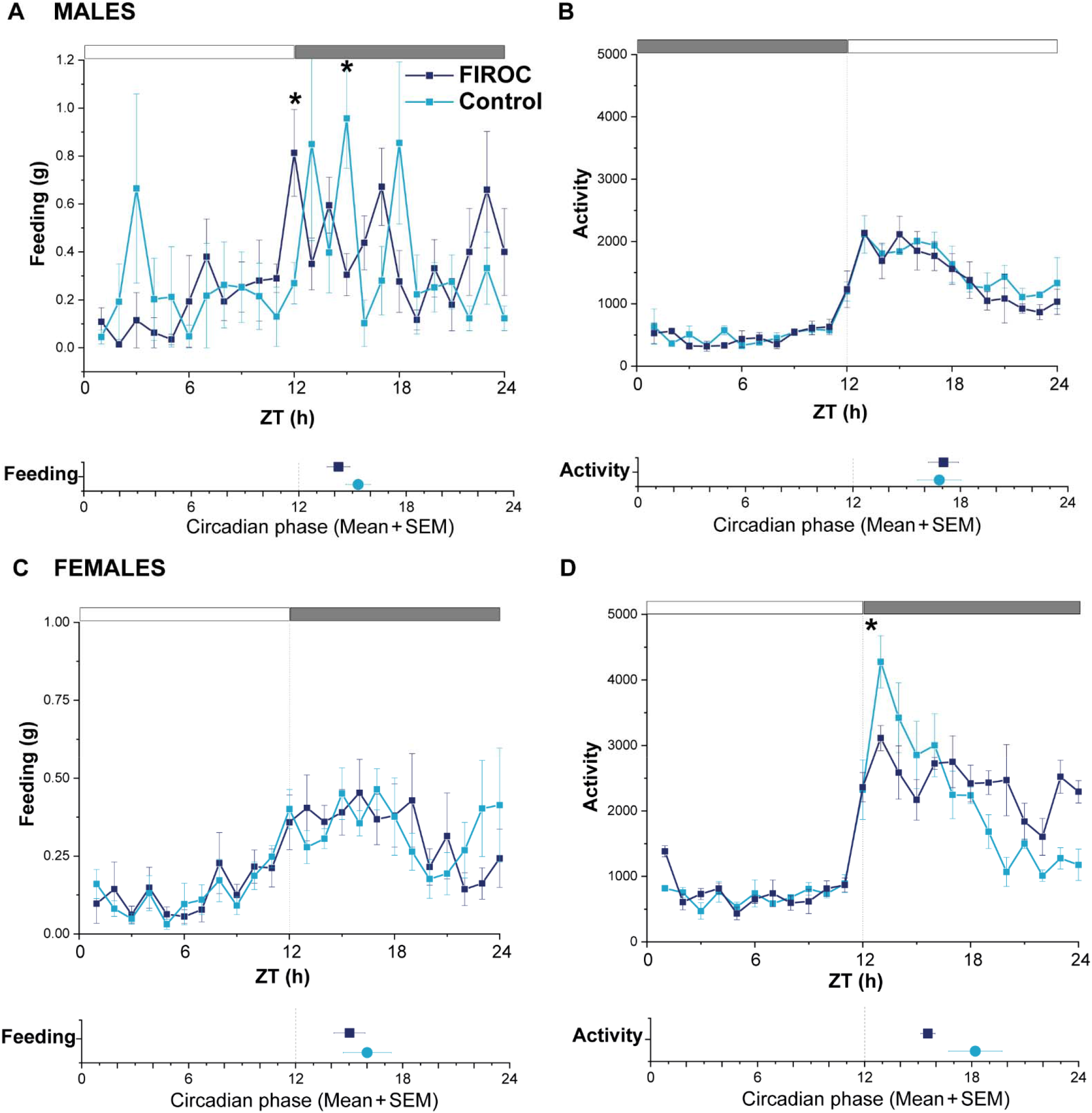
Circadian behavior is disturbed in Firoc mice. **A-D,** Firoc (n= 7-9), and littermate control (n=7) male (A, B) and female (C, D) mice were housed in Phenomaster cages for 14 days. Food intake (A, C) and activity (B, D) were registered daily every 1h for 24h. The circadian phase obtained for each parameter using the Metacycle algorithm is shown at the bottom of each panel. Data are the mean ± SEM for the last 5 days at each time point (two-sample t-test, *p<0.05).

### IGF-I modulates metabolic homeostasis through orexin neurons

We next evaluated whether the circadian misalignment and loss of glucose rhythm observed in Firoc mice impacted on metabolic homeostasis. Indeed, Firoc males developed hyperglycemia as they aged, while females did not show differences in blood glucose levels (Figure 3A). However, Firoc females, but not males, showed significantly increased weight with age (Figure 3B) and accumulated significantly more visceral and pelvic fat tissue than control littermates (Figure 3C, D).

**Figure 3:**
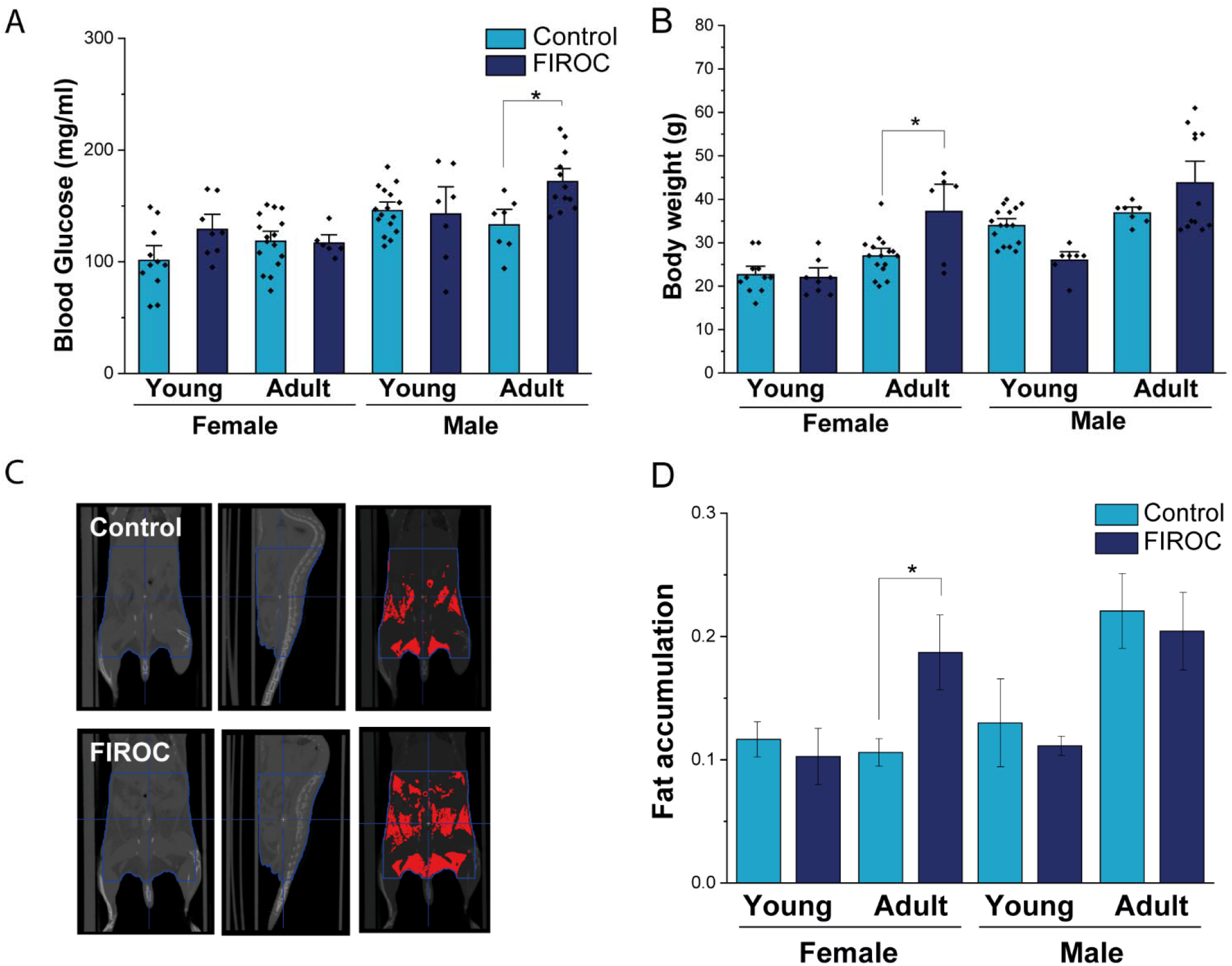
Sex-dependent metabolic disturbances in Firoc mice. **A,** Basal blood glucose levels after 6h fasting were significantly increased in males, but not in female Firoc mice, in an age-dependent fashion. **B,** Body weight was significantly greater in female, but not male Firoc mice, in an age-dependent manner. **C,** Representative CT scans illustrating visceral and pelvic fat deposits (red masses in right panel) in female Firoc and littermates. Whole body ventral (left panels) and lateral (central panels) views are depicted. **D,** Body adiposity was significantly increased in adult Firoc females, but not in males (One-way ANOVA; *p<0.05).

In agreement with the above results, adult, but not young (Figure S1A,B) Firoc males displayed glucose intolerance (Figure 4A). Their insulin sensitivity was not affected (Figure 4B). Neither adult (Figure 4C,D) nor young Figure S1C,D) Firoc females show altered glucose or insulin sensitivity. Nevertheless, since adult Firoc females showed increased body weight, we analyzed their use of alternative metabolic pathways. Adult Firoc females showed increased responsiveness to pyruvate (Figure 4E), an alternative fuel in liver gluconeogenesis, as significantly larger glucose levels were seen after pyruvate load (Figure 4H).

**Figure 4:**
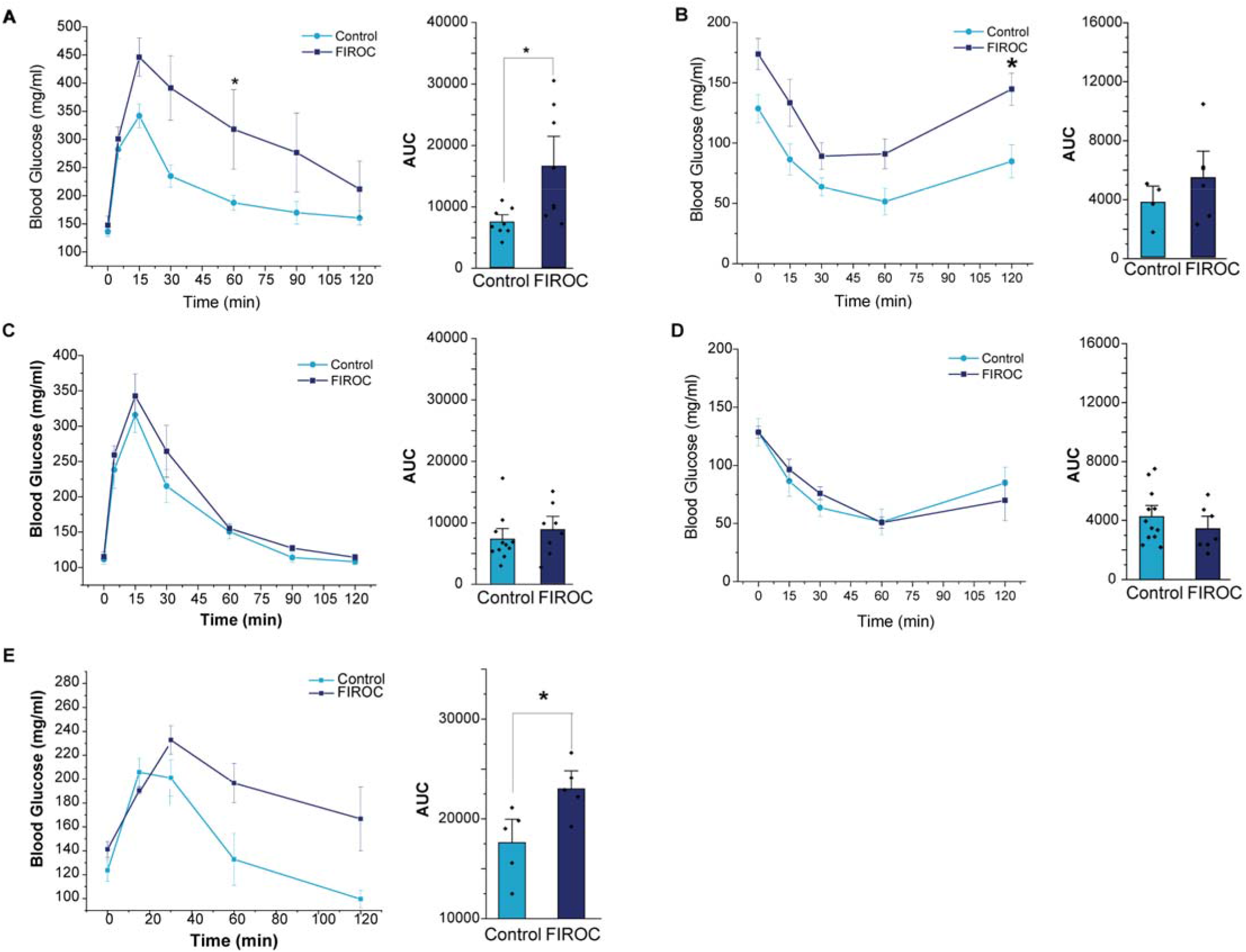
Peripheral glucose homeostasis is altered in male but not in female Firoc mice. **A,C,** After glucose load, male Firoc mice (A), but not females (C), show higher glucose levels throughout the 2 h test. This is reflected in a significantly greater area under the curve (AUC) of blood glucose levels (histograms on the right). **B,D**, Glucose levels after systemic insulin administration remained within normal values in both male (B) and female (D) Firoc mice. AUC histograms are shown in right panels. **E,** Glucose levels were significantly increased in Firoc females after pyruvate administration as reflected in AUC values (right histograms). Two way ANOVA and Tukeys test (*p<0.05).

### IGF-I regulates orexin to entrain metabolic clock with circadian rhythm

Synchronization of hepatic and body circadian clocks is partially mediated by Krüppel-Jacob transcription factors (KLF)-10 and −15 through regulating the expression of liver gluconeogenic enzymes^33^. Hence, we analyzed their liver expression after intracerebroventricular (icv) administration of IGF-I (1 μg) at ZT10 and found increased levels of KLF-10 in control, but not Firoc mice (Figure 5A) at ZT12. Rather, Firoc mice show increased KLF-15 expression in response to IGF-I (Figure 5B).

**Figure 5:**
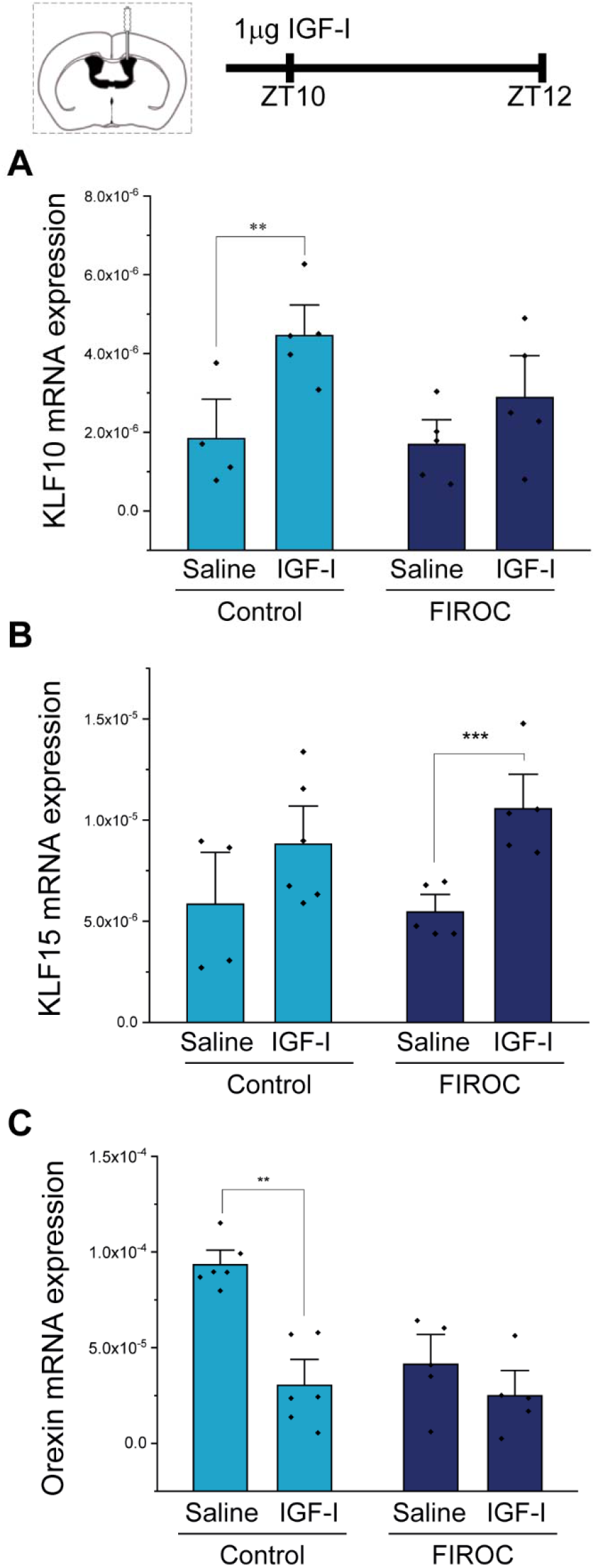
Liver KLF10 expression is regulated by IGF-I signaling in orexin neurons. Cannulae were placed icv in Control (n=10) and Firoc mice (n=10). After mice recovered from surgery, 1ug of IGF-I or saline were icv injected at ZT10 and mice were sacrificed 2 hours later. Transcriptional changes in the liver (**A, B**) and in the hypothalamus **(C)** were analyzed by real-time **PCR**. IGF-I icv administration significantly increased KLF-10 expression (A) in the liver in control, but not in Firoc mice. Conversely, IGF-I increased KLF-15 (B) in Firoc but not in control mice. In the hypothalamus, IGF-I decreased orexin transcription (C) in control, but not Firoc mice (t-test, *p<0.05).

Based on the above observations we determined whether IGF-I could entrain the hepatic clock with the circadian cycle by regulating hypothalamic orexin expression. We administered icv IGF-I at ZT10 (end of the inactive phase) and measured hypothalamic levels of orexin mRNA at ZT12 (beginning of the active phase). We found decreased orexin expression in control, but not in Firoc mice after IGF-I (Figure 5C). Similar results were obtained in primary hypothalamic neuronal cultures, where IGF-I (1 nM) significantly decreased orexin mRNA levels (Figure S2).

Given that IGF-I appears to be an entrainment signal modulating orexin to synchronize the circadian and hepatic cycles, we determined whether these cycles were uncoupled in Firoc mice. We kept Firoc mice under constant darkness -to remove light entrainment and restricted food availability to 12h/day (between ZT0-ZT12). As expected, 5 days later mice were able to synchronize their activity with food availability (Figure 6A). However, control mice mainly had meals during dusk, while Firoc mice did not distinguish between dusk and dawn (Figure 6B). In a second experiment, we measured food anticipatory activity (FAA), using free running-wheels as a proxy of the total activity of the animals, as a behavioral readout of the metabolic cycle. After 1 week of habituation in normal light conditions and *ad libitum* feeding, the light was removed and feeding time was restricted to 6h per day from ZT6 to ZT12. Mice get adapted to the new feeding schedule in a few days, showing increased activity 3 hours before food presentation (Figure 6C), while total food intake was not different between groups (data not shown). After 5 days, FAA in Firoc mice was higher than in littermates (Figure 6D).

**Figure 6:**
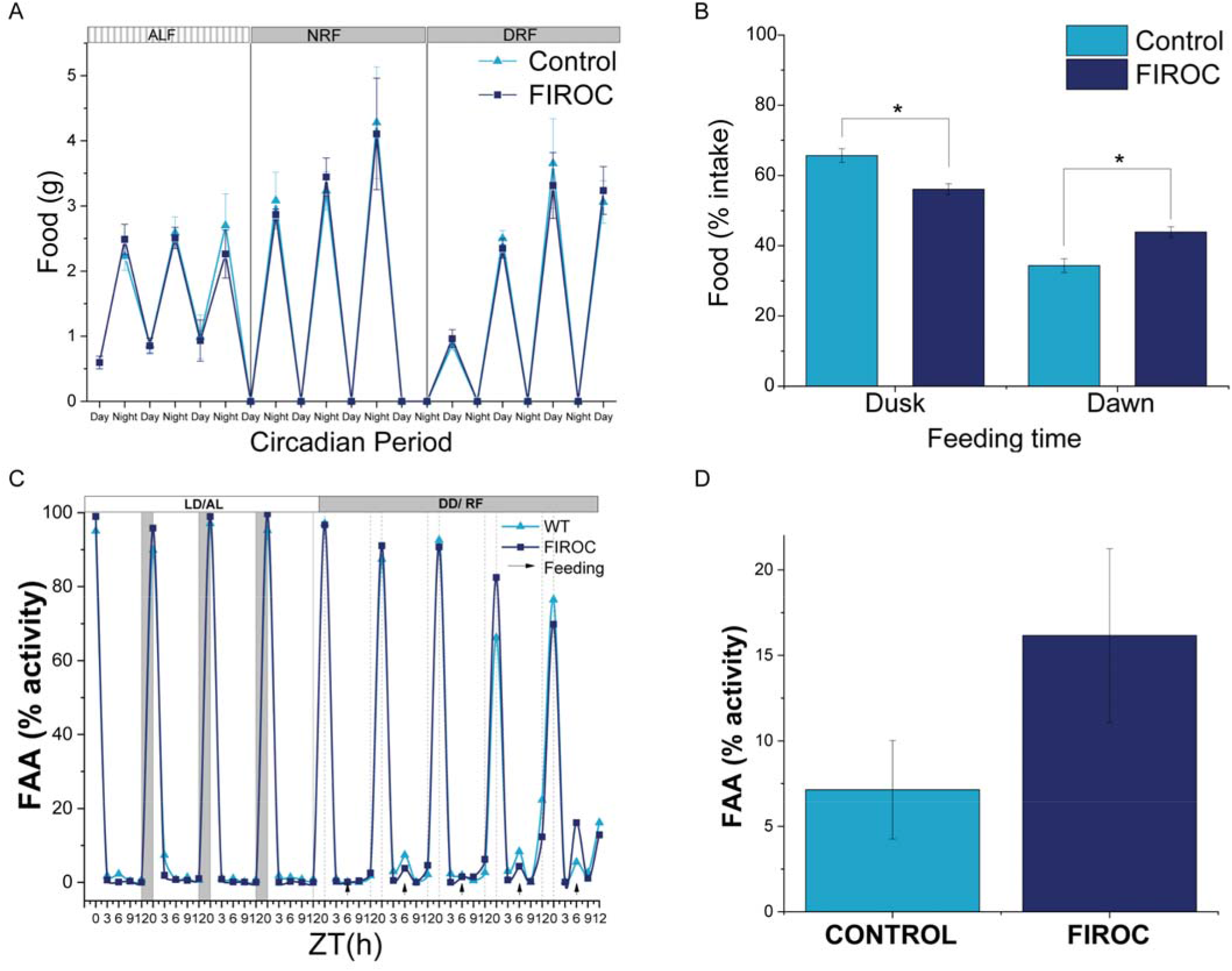
Circadian and food clock alignment is impaired in Firoc mice. **A**, Both Firoc and control mice adapted to changes in light/food presentation. Mice were housed individually for 1 week under a regular light/dark cycle and *ad libitum* food regime (ALF). Then, lights were removed and food was restricted to the subjective night (NRF) for 4 days. After that, food availability was moved to the subjective day (DRF). Both groups spent about 4 days to become habituated to these new conditions and ate a similar amount of food as those observed during the ALF phase. **B,** However, analysis of the amount of food taken during the first (Dusk) and second 6 hours (Dawn), showed that control mice mainly had meals during dusk, as expected, while Firoc mice did not distinguish between dusk and dawn (t-test, *p<0.05). **C,** Food anticipatory activity (FAA) was measured under constant darkness after 5 days of adaptation when both control and Firoc mice ingested the same amount of food. **D,** Firoc mice showed an increased FAA 3h before food presentation as compared to control mice (n=3 per group).

## Discussion

These results suggest that impaired IGF-I signaling onto orexin neurons alters the temporal relationship between feeding and/or activity with light cycles, suggesting that the synchronization between metabolic and circadian rhythms is altered. This confirms and extends previous observations that IGF-I entrains feeding and circadian rhythms^16^, pointing to orexin neurons as a site of food/light entrainment by IGF-I. Indeed, hypothalamic orexin neurons regulate sleep/wake rhythms^12,27,34^ and feeding behavior^32^, and have been described as food-entrained oscilators^35^, as orexin knockout mice display reduced food anticipatory activity^36^. Conceivably, the circadian activity of IGF-I ^37^ may signal orexin neurons to help maintain metabolic homeostasis since blockage of IGF-I favors the development of metabolic disorders.

IGF-I administration at the end of the resting phase reduced expression of orexin at the beginning of the active phase, a time where orexin expression surges^29^. A direct inhibitory effect of IGF-I was shown in cultured hypothalamic neurons. Certainly, the circadian activity of IGF-I shows a trough at the end of the resting phase, before the surge in orexin levels^38^. Together with the observation that circadian variation of hypothalamic orexin is disturbed in Firoc mice, these data suggest that IGF-I signaling onto orexin neurons is important for the regulation of orexin expression, as previously suggested^27^.

IGF-I also regulated the expression of KLF10 in the liver. KLF10 belongs to the Krüpel-like transcription factor family that orchestrate systemic metabolism through local and multi-organ transcriptional regulation of all types of macronutrients^38^. KLF10, as well as KLF15, are known circadian and metabolic entrainment transcription factors and their expression in the liver follows a circadian rhythm regulated by clock genes and the light cycle^33^. KLF10 regulates hepatic glucose production by directly regulating the expression of gluconeogenic enzymes^33^. Thus, KLF10 knockout mice develop alterations in glucose metabolism in a sex-dependent manner, with hyperglycemia in males and obesity in females^33^, reminiscent of the changes observed in Firoc mice. These results suggest that orexin neurons may participate in the synchronization of the circadian clock with hepatic metabolism by regulating KLF-10 expression at the beginning of the active phase. This synchronization would be mediated by IGF-I, as Firoc mice do not increase KLF-10 in response to this hormone. An intriguing observation is that KLF-15 increases in Firoc mice in response to IGF-I, suggesting the existence of compensatory, orexin-independent regulation of hepatic function by IGF-I.

IGF-I entrainment of metabolic and circadian rhythms through orexin neurons regulates circadian glucose rhythms in a sex-specific manner. Alteration of this rhythm could be either the cause or the consequence of the mild sex-dependent disruption of circadian behavior observed in Firoc mice, emphasizing the sex-dependent actions of both IGF-I and orexin^28,39^. Daily blood glucose fluctuations are regulated by the SCN through its glutamatergic and GABAergic projections to the PVN^40,41^ and to the LH^24^, especially to orexin neurons, and their respective para- and sympathetic innervations to the liver. This central glucose regulation promotes a peak in blood glucose synchronized with the beginning of the active phase^40,42^. The role of orexin neurons in circadian control of hepatic glucose production is supported by the observation that activation of orexin neurons promotes glucose release^32^, and by the fact that orexin knockout mice lack a glucose circadian rhythm. The latter is due to an impaired liver innervation and can be restored by orexin administration at the end of the resting phase^30^. All these results reinforce the relevance of IGF-I as a peripheral signal for central glucose homeostasis in conjunction with circadian rhythms.

Circadian and metabolic cycles are desynchronized in Firoc mice. The circadian cycle driven by light and the SCN is known as the circadian “pacemaker” and synchronizes peripheral clocks, including the hepatic clock, through the HPA axis^9^. In turn, the liver has its own circadian clock that prevails when the SCN is uncoupled. We now provide evidence that IGF-I tunes orexin neurons to modulate the metabolic or food clock, following the food anticipatory activity. The anatomical region involved in this phenomenon remains unclear, but it has been demonstrated to be SCN-independent^43–45^ and dependent on other neuronal populations from the hypothalamus and perifornical area including orexin neurons since orexin knockout mice show reduced FAA^35,36^. It seems that in Firoc mice the food clock predominates over the circadian clock, as they show enhanced FAA. This unbalance could account for altered glucose homeostasis that may antecede metabolic alterations observed in Firoc mice.

Overall, we observed a new aspect of orexin activity in circadian and metabolic alignment where IGF-I is involved. This alignment integrates circadian inputs coming from the SCN with metabolic inputs coming from the periphery. Hence, alterations in orexin function could be a consequence of IGF-I dysregulation due to aging, diet, or pathology. These results indicate that IGF-I regulates orexin neurons, and consequently circadian rhythm and metabolism, extending and confirming our previous observations about the central actions of IGF-I on mood, metabolism, and wellness^27,46^.

## Material and Methods

### Animals

Adult female and male C57BL/6J mice (Harlan Laboratories, Spain) and Cre/Lox mice lacking functional IGF-I receptors in orexin neurons (Floxed IGF-IR/Orexin Cre: Firoc mice) were used. Littermates were used as controls. The estrous cycle of the female mice was not controlled. Experiments were done during the light phase, except when indicated, Firoc mice were obtained as described in detail elsewhere^27^ by crossing Orexin-Cre mice (a kind gift of T Sakurai, Tsukuba Univ, Japan) with IGF-IR^f/f^ mice (B6, 129 background; Jackson Labs; stock number: 012251; exon 3 floxed). Animals were housed in standard cages (48 × 26 cm^2^, 5 per cage), and kept in a room with controlled temperature (22°C) under a 12-12h light-dark cycle. Mice were fed with a pellet rodent diet and water *ad libitum*. Animal procedures followed European guidelines (2010/63, European Council Directives) and were approved by the local Bioethics Committee (Government of the Community of Madrid).

### Metabolic cages

Firoc (3-4 months; n=6-8, both sexes) and control littermates (n= 6-7 both sexes) were housed in metabolic cages. Movements, food intake and drinking, energy expenditure (EE), respiratory exchange ratio (RER), and consumed volume of O2 (VO2) were obtained. Data was collected over 24h at 1h intervals for five days. Circadian rhythms were analyzed using an algorithm (Metacycle)^47^ to obtain the Period, Amplitude, Phase, and Mesor values for each metabolic parameter.

### Behavioral circadian test

In a first experiment, animals were maintained for 1 week under normal (12:12) 1ight:dark regimen and *ad libitum* feeding, adding measured food pellets (5g) into the cage at ZT0 and ZT12, and food consumption was weighed. After 1 week, the light was switched off at ZT0, mice were fed *ad libitum,* and food weighed at the end of each period. Finally, 1 week later, food was only supplied from ZT0 to ZT12, their corresponding inactive phase, and food was weighed at ZT12. Five days later, food was weighed at ZT6 and ZT12. In a second experiment to analyze spontaneous activity, mice were housed individually in cages with a vertical running wheel and an automatic wheel counter. Mice were kept under normal conditions for 1 week and running was scored at ZT3, ZT6, ZT9, and ZT12. At the same time, food consumption was measured as described. After 1 week, lights were switched off at ZT0, and food was provided only from ZT6 to ZT12. The activity was recorded at ZT3, 6, 9, and 12, and food consumption was measured at ZT9 and ZT12. Spontaneous activity was registered as wheel counts between ZT3 and ZT6, the time before food supply, and it was only taken into account after 5 days of light and feeding conditioning.

### Glucose, insulin, and pyruvate tolerance tests

For the Glucose tolerance test (GTT), 6h fasted mice were injected with a glucose bolus (2g/kg) intraperitoneally (IP) and blood glucose levels were measured at 0, 5, 15, 30, 90, 120 min after injection using a blood glucometer (Glucomen areo, A. Menarini Diagnostics, Italy). For the Insulin Tolerance test, mice were fasted for 6h and then 1U/kg human Insulin (Actrapid Penfill, Novo Nordisk A/S, Denmark), was IP injected. Glucose levels were measured at 15, 30, 90, 120 min after injection. For the Pyruvate tolerance test, mice were fasted for 16h, then 2g/kg Sodium pyruvate was IP injected, and glucose tolerance test was measured at 15, 30, 90, and 120 minutes after injection. Glucose levels were determined using Glucomen aero 2K glucometer (A. Menarini diagnostics).

### Computed Tomography

Body fat content analysis was carried out as described^48,49^. Briefly, mice were anesthetized by Isofluorane (2.5% flow rate) and kept under at 2.5% via a nose-cone setup for imaging. Animals were positioned prone with limbs lateral from the torso for a uniform CT acquisition. Image acquisitions were performed using the Albira II SPECT/CT (Bruker BioSpin PCI) with 600 projections. The X-ray source was set to a current of 400 μA and voltage of 45 kVp. Images were reconstructed using the FBP (Filtered Back Projection) algorithm via the Albira Suite 5.0 Reconstructor using “Standard” parameters. These combined acquisition and reconstruction settings produce a final image with 125 μm isotropic voxels. Images were segmented using POMD v3.3 software according to tissue density-first for total volume, and then, for fat volume segmentation, values between −500 to −100 Housfield units were considered.

### Neuronal cultures

Brains were dissected from E15-16 mouse embryos. Meninges and blood vessels were removed, and the hypothalamus dissected, minced with scissors and dissociated with papain at 37 °C/2h in agitation. Neuronal cultures were grown in Neurobasal medium supplemented with B27 (Gibco) and Glutamax (Gibco) in 6-well plates coated with poly-L-lysine. Ten days later, cultures were washed with PBS, B27-free Neurobasal added for 3 hours, and then treated with IGF-I (Pre-Protech, USA) at 1nM for 15h. Thereafter, plates were washed with PBS, and Trizol added for RNA extraction.

### IGF-I injection

Guide cannulae (2.5mm in length, Bilaney) were implanted by stereotactic surgery at coordinates 0.2mm A/P, 0.9mm M/L, and 2.5mm in-depth from the skull. After 1 week of recovery, mice were fasted for 24h to promote orexin activity ^30^. At ZT10, mice were lightly anesthetized to remove the dummy cannula and place the internal cannula. Once mice were awake in the cage (2-3 min), 1 μl containing 1 μg of IGF-I (Pre-Protech, USA) in saline solution was injected at 1μl/min. The internal cannula remained placed for 1 extra minute before removal. Mice were kept in the cages for 2 hours until ZT12 and sacrificed, cardially perfused and hypothalamus and liver samples collected and stored at −80°C until use.

### RNA isolation and real-time PCR

Tissue RNA was extracted with Trizol (Life Technologies, USA), as described elsewhere^50^. cDNA was synthesized from 1μg of RNA of each sample following the manufacturer’s instructions (High Capacity cDNA Reverse Transcription Kit; Applied Biosystems). Fast Real-time qPCR was performed using the SYBR Green method (Fast SYBR Green Master Mix, Applied Biosystems) with the QuantStudio 3 Real-Time PCR System (Applied Biosystems). Relative mRNA expression was determined by the 2^-ΔΔCT^ method (Pfaffl,2001), and normalized to ribosomal 18S mRNA levels.

### Statistics

Statistical analyses were performed with OriginPro (OriginLab Corp., Northhampton, USA). F-Test or Levene’s test was conducted for variance test previous to Student’s t-test, when comparing two groups, or either 1-way or 2-way ANOVA (for more than 2 groups) followed by Tukey’s multiple comparison test. All results are shown as mean ± standard error (SEM) and significant values as: *p<0.05; **p<0.01; and ***p<0.001.

---

**Table S1: Algorithmic analysis of metabolic circadian rhythms.**

Data obtained from Phenomaster metabolic cages in male (n=8 per genotype group) and female (n=8 per genotype group) Firoc and littermate mice were analyzed using Metacycle algorithm. Mean and SEM of the period, phase, and amplitude parameters from feeding, water intake, respiratory exchange ratio (RER), energy expenditure (EE), activity on X and XY axis and volume of oxygen consumed (VO2) were obtained.

**Figure S1: Peripheral glucose homeostasis in young Firoc mice is unaltered.** Glucose levels remain in young male (A,B) and female (C,D) Firoc mice within control range after glucose (A,C) or insulin (B,D) load. AUC histograms are shown in right panels.

**Figure S2: IGF-I decreasess orexin expression in hypothalamic neurons.**

Hypothalamic neurons from E13 mice embryos (n=5) were cultured for 10 days. Cells were treated with IGF-I or saline for 15h. IGF-I significantly decreased orexin mRNA transcription as compared to saline treated cells (mean fold change ± SEM, t-test, *p < 0.05).

