## Supplementary figures and images for "Insulin-Like Growth Factor I Couples Metabolism With Circadian Activity Through Hypothalamic Orexin Neurons"

### Supplemetary data

#### Slide 1
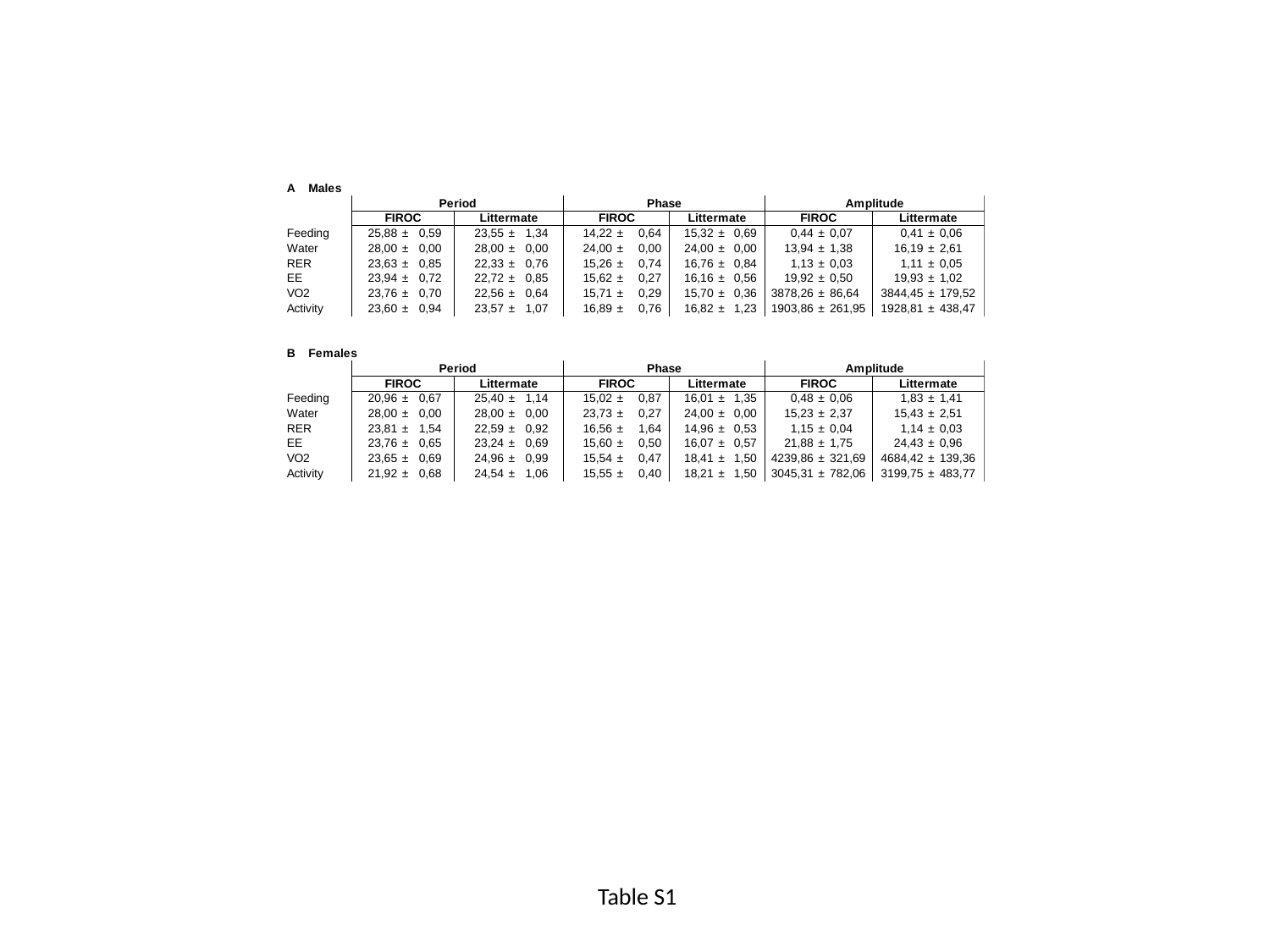

Table S1

#### Slide 2
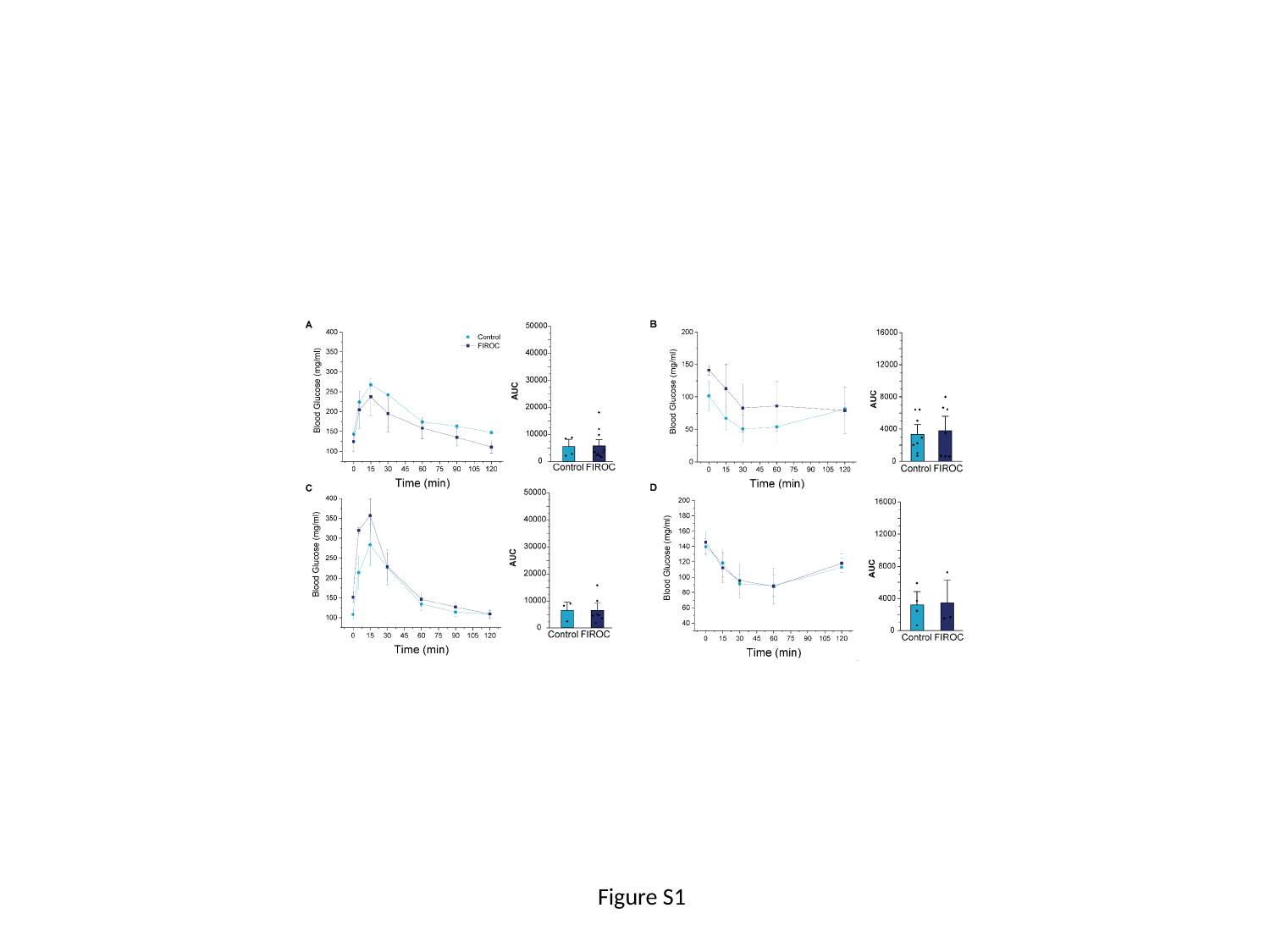

Figure S1

#### Slide 3
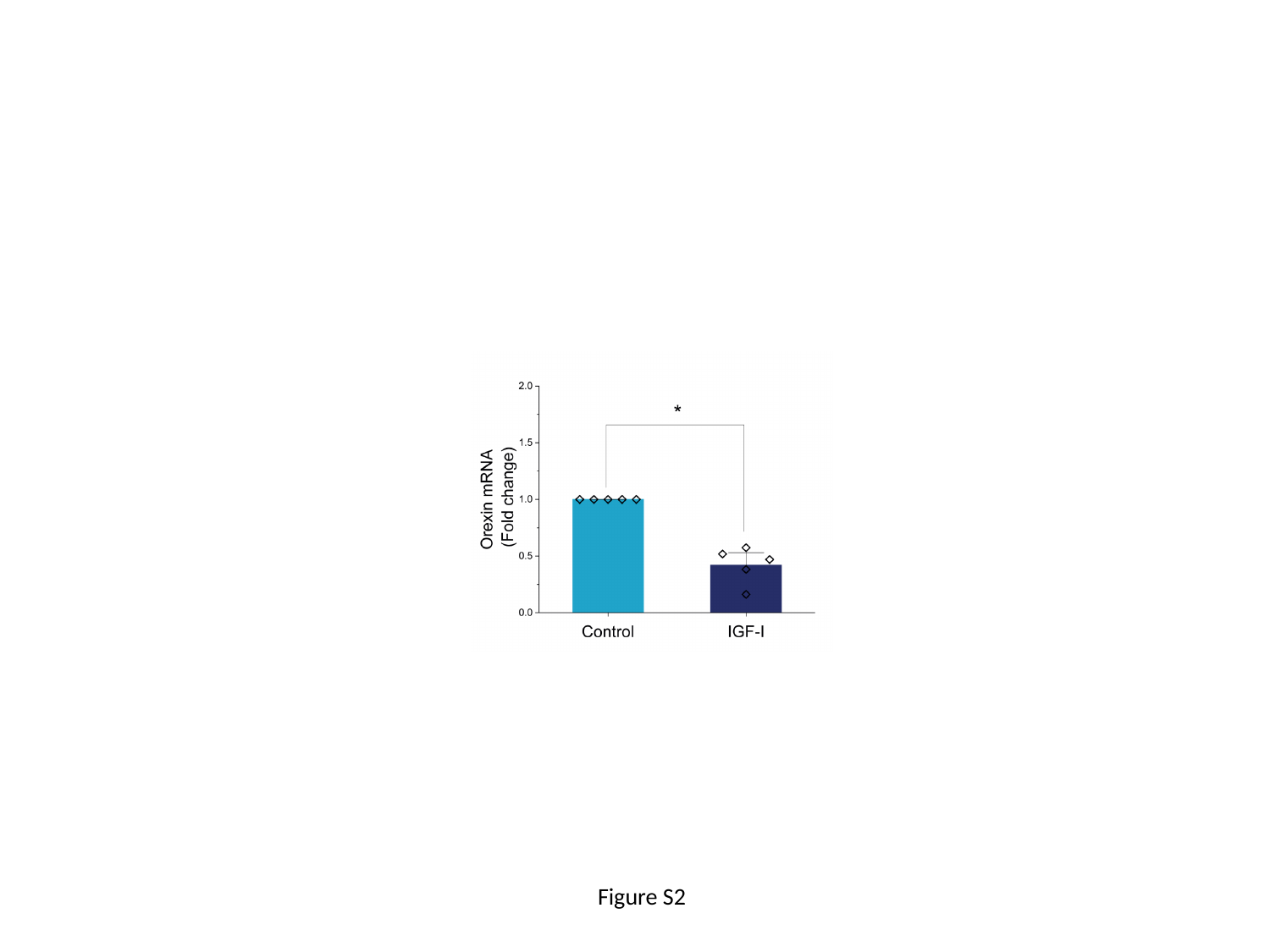

Figure S2
